# Diversification and coexistence amid widespread and persistent gene flow in Andean trees

**DOI:** 10.1101/2025.09.22.677938

**Authors:** Ellen J. Quinlan, James B. Pease, Jhonatan Sallo Bravo, Alfredo F. Fuentes, William Farfan-Rios, Miles R. Silman

## Abstract

- Hybridization and introgression are increasingly recognized as important in plant evolution, yet their prevalence and consequences remain poorly understood in tropical Andean trees, where species coexist at low densities and turn over rapidly across elevation gradients.
- Here, we investigate whether species boundaries are maintained among congeneric Andean *Prunus* distributed across the Andes-Amazon elevational gradient by reconstructing phylogenetic relationships, divergence times, and patterns of historical and recent gene flow.
- We find that Andean *Prunus* diversified during the Miocene, likely in response to novel habitat opportunities resulting from mountain uplift, combined with allopatric processes. Despite this evolutionary divergence, these species show extensive ancient introgression and recent admixture, indicating that species boundaries remain permeable across both ecological and phylogenetic distance.
- Together, these results suggest that Andean *Prunus* comprise distinct but evolutionarily connected lineages that retain morphological identity while potentially functioning as a long-lived syngameon, with implications for understanding the origin and maintenance of tropical tree diversity and future response to global change.

## INTRODUCTION

Coexistence theories of ecology generally assume species are discrete evolutionary units (e.g. Chesson 2000, Hubbell 2005), though interspecific gene flow is known to have been common throughout the evolutionary history of plants (reviewed by Stull et al. 2023). No system lends itself to tests of these theories better than species-rich tropical forests. Until recently, hybridization was considered an extremely rare occurrence in tropical trees, as hybrids were thought to be poor competitors in diverse systems and few individuals with intermediate morphologies have been identified (e.g. Ashton 1969; Ehrendorfer 1970; Gentry 1982). Mounting genomic evidence suggests hybridization has been widespread throughout the diversification of the Neotropical flora (reviewed by Schley et al. 2022) and may be an important evolutionary mechanism for maintaining high tree diversity with low population densities (Cannon and Lerdau 2015, Cervantes et al. 2025, Quinlan et al. 2025). These findings collectively point to the importance of reevaluating the impact of hybridization in our understanding of species diversity and coexistence theories in tropical ecosystems.

The tendency for inter-specific gene flow and hybridization to occur amongst closely related, sympatric trees has been widely documented in temperate zones (De La Torre et al. 2014, Tsuda et al. 2017, Cavender-Bares et al. 2018, Chhatre et al. 2018, Menon et al. 2018), yet species cohesion and morphological similarity is often maintained (Grant 1971; Gailing and Curtu 2014; Cannon and Petit 2020). This may be partially due to assortative (non-random) mating and variable mating behaviors (selfing, conspecific outcrossing, and interspecific gene flow), determined by population densities and pollinator abundance (Hamrick and Murawski 1990; Ward et al. 2005; Cannon and Lerdau 2015). When present, conspecific pollen can outcompete heterospecific pollen, reinforcing assortative mating (Williams Jr. et al. 1999). However, incomplete reproductive isolation and the ability to accept heterospecific pollen may become adaptively beneficial when conspecific pollen is rare, as the benefit of avoiding local extinction overrides potential negative consequences of hybridization such as reduced hybrid fitness (Barton 2001, Wu 2001, Cannon and Lerdau 2015). Hybridization with introgression may be particularly beneficial in times of rapid environmental change, as it can produce novel gene combinations within a single generation, increasing adaptive potential and population resilience (Anderson and Stebbins 1954, Baskett and Gomulkiewicz 2011, Suarez-Gonzalez 2018, Brauer et al. 2023, Kulmuni et al. 2024). Such density-dependent mating has been reported in oaks (Lepais et al. 2009) and may be particularly relevant to tropical trees which maintain extremely low local population densities (Pires et al. 1953, Pitman et al. 1999, ter Steege et al. 2013).

Most tropical forest trees that have been studied are highly outcrossed (Bawa et al. 1985a, b, Hamrick and Murawski 1990, Ward et al. 2005), though individuals retain variable capacities for different mating behaviors, even within populations and across flowering events (Murawski and Hamrick 1991, Nason and Hamrick 1997). Self-fertilization has been shown to be inversely correlated with population density (Hamrick and Murawski 1990, Ward et al. 2005), but few studies have assessed the extent of inter-specific gene flow. Those that have (e.g. Schley et al. 2020 – *Brownea*, Larson et al. 2021 – *Eschweilera,* Schley et al. 2025 – *Inga*), detected widespread admixture. Evolutionary distance may be one important constraint on interspecific gene flow among congeners in sympatry, as hybridization is thought to decrease with increasing divergence due to the accumulation of pre- and postzygotic barriers through time (Mendelson et al. 2004, Levin 2012, Comeault and Matute 2018). Linan et al. (2021) found evidence of this in the Mascarene Island *Diospyros* clade, suggesting the processes by which species diversify and assemble into communities may have lasting effects on their eco-evolutionary dynamics. For example, adaptive radiations across gradients typically result in stepwise diversification patterns with sister taxa occurring closest in niche proximity (Simpson 1953, Schluter 2000, Stroud and Losos 2016), whereas communities assembled predominantly via the immigration and sorting of pre-adapted clades demonstrate greater phylogenetic distance among co-occurring taxa (Donoghue 2008, Dexter et al. 2017, Linan et al. 2021).

In tropical montane forests, genera commonly comprise many rare species that turn over rapidly with elevation (Terborgh 1971, Gentry 1982). Turnover in the abiotic environment and biotic interactions leads to narrow, shoestring-like distributions and the packing of many closely related species across elevation gradients (e.g. Janzen 1967, Jankowski et al. 2009, Hillyer and Silman 2010, Freeman et al. 2022). In the tropical Andes, this is consistent across taxa and is especially true for forest trees (e.g. Terborgh 1977, Meier et al. 2010, Jankowski et al. 2012). While many tree species have elevation ranges of <500 m, the distributions of entire genera are often much broader, with congeneric species occurring in partial sympatry, especially at range margins (Feeley et al. 2011, Griffiths et al. 2021). At the genus level, immigration and sorting of pre-adapted clades has been shown to be the primary mechanism structuring tree diversity across elevation gradients in the Andes (Griffiths et al. 2021, Segovia et al. 2020, Linan et al. 2021). However, many examples of adaptive diversification have been found within individual plant clades (e.g. Nϋrk et al. 2013, Lagomarsino et al. 2016, Nevado et al. 2016, among many others). Still, few studies have explored the evolutionary relationships among congeners within individual tree clades that co-occur across the gradient, and characterizing these evolutionary relationships is essential for understanding potential gene flow and its ecological implications.

*Prunus* L. (Rosaceae) is one such genus that exhibits high species turnover with elevation in the Andes (Figure 1). Globally, it has diversified into a range of environments, and though typically considered a temperate group (Wen et al. 2008, Chin et al. 2014), a recent revision found its highest diversity occurs in the tropics and primarily the Neotropics (∼228 Neotropical species; Pérez-Zabala 2022). The Andes contain the highest number of species, with 41 total taxa including 39 endemics identified from the Central Andes alone. A total of 33 taxa, including 10 endemics are known from the Amazon, Guianas and northern South America (Pérez-Zabala 2022). *Prunus* is one of the only tree genera with species spanning the entire Andes-Amazon gradient, from treeline (∼3800 m) into the lowlands (<500 m), though individual species have more narrow ranges, typically 500–1000 m or less (e.g. Achá 2013). It is a keystone clade, whose fruits are important in the diets of frugivorous birds, primates, Andean bears, and humans (Achá 2013). The genus also contains many globally important crop species (peach, plum, almond, cherries) and others used as ornamentals, timber, and medicine (Andro and Riffaud 1995, Bortiri et al. 2001, Lee and Wen 2001, Wen et al. 2008).

**Figure 1.**
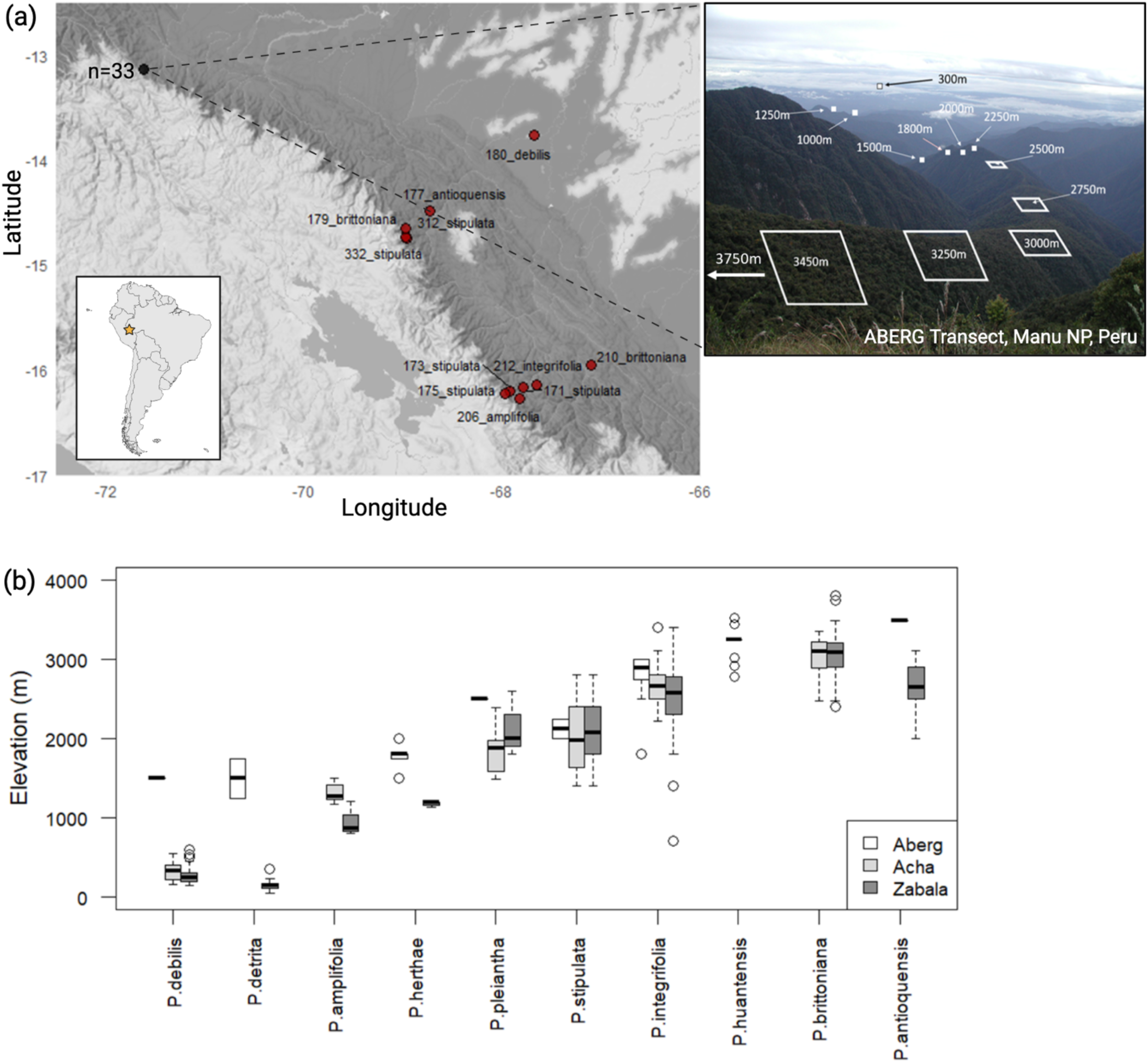
(a) Map of samples included in this study. Black dot indicates the location of the ABERG transect in Manu National Park, Peru where most samples (n=33) were collected, also shown in picture to the right. Red points indicate locations of additional samples included in the study. Further details on individual samples provided in Table S1. (b) Box plots showing elevational distributions of the in-group Andean species across three different sources. “Aberg” includes all known individuals along the ABERG transect >10 dbh over 20 years. “Acha” includes all Bolivian material reviewed by Achá (2013) and “Zabala” includes all material reviewed by Pérez-Zabala (2022) from Venezuela to Bolivia. Pérez-Zabala considered *P. huantensis* to be *P. brittoniana* so those samples are grouped together on the figure under *P. brittoniana*. Figure does not include new “varieties” of species proposed by either Achá or Pérez-Zabala.

The genus *Prunus* is divided into three major clades that correspond to inflorescence type: solitary, corymbose, and racemose (Rehder 1940, Su et al. 2023). All species endemic to the Americas and tropical regions are racemose (Pérez-Zabala 2022). Though historically understudied, the exclusively South American species studied to date form a strongly supported clade nested within the tropical racemose group (Hodel et al. 2023—11species, Hyb-Seq and full plastome sequencing; Pérez-Zabala 2022—30 species, PCR). Studies have placed the origin of the genus in either North America (Chin et al. 2014) or East Asia (Hodel et al. 2023) during the Paleocene (ca. 68–55 Ma), with the three major clades diverging by the early-mid Eocene as part of the Boreotropical flora (Chin et al. 2014, Hodel et al. 2023, Su et al. 2023). Divergence in the tropical Racemose group occurred in several regions including Asia, Africa, North America, and Europe ca. 55–40 Ma (Hodel et al. 2023). Chin et al. (2014) hypothesized migration into South America occurred via the dispersal of a North American relative ca. 34 Ma through Central America and the Caribbean. Hodel et al. (2023) found evidence of a long period of stasis in the Neotropical lineages followed by rapid diversification during the Miocene.

While most species in the solitary-flower and corymbose groups are diploids (2n=2x=16), the racemose lineage is known for higher ploidy levels, with allopolyploidy and/or hybridization thought to have driven early diversification in the genus (Chin et al. 2014, Zhao et al. 2016, Hodel et al. 2021, Su et al. 2023). Hodel et al. (2023) concluded that the majority, if not all racemose species are polyploid, and all tropical species studied to date have been tetraploid (2n=4x=32). The two exceptions being *P. laurocerasus* (2n=22x=176) and *P. gongshanensis* (2n=8x=64)—species native to old world Europe and Southeast Asia (Hodel et al. 2023, Xie et al. 2025). Domesticated *Prunus* (particularly plums) readily hybridize in agricultural settings (Cici and Acker 2010), resulting in the creation of some crops (e.g. *P. armeniaca* L. × *P. cerasifera* Ehrh. – ‘black apricots,’ *P. salicina* × *P. armeniaca* Lindl. – plumcots, pluots, and apriums; Guerrero et al. 2022), as well as among cultivated varieties and wild conspecifics (*P. yedoensis* - Baek et al. 2018). Species in this genus are highly outcrossing and typically self-incompatible with generalist insect pollinators (Cici and Acker 2010), though we are unaware of any study that has specifically looked at either breeding systems or pollination in Neotropical *Prunus*.

Here, we investigate the extent to which species boundaries are maintained among congeneric Andean trees, across elevation gradients and through time. Specifically, we ask: (1) what are the evolutionary relationships among species that turnover with elevation, (2) are these species are genetically isolated and (3) is genetic isolation related to evolutionary or ecological distance, with ecological distance defined as elevational range overlap. To do this, we reconstructed the phylogenetic relationships among sympatric *Prunus* species, estimated the timing of divergence, and tested for historical introgression and recent admixture using RADseq data. The results of these analyses expand current understanding of the origin and maintenance of tropical Andean tree diversity and provide important context for predicting how these species may respond to rapid environmental change.

## MATERIALS AND METHODS

### Study site and sample collection

Sampling for this study primarily focused on species which co-occur across the Andes Biodiversity and Ecosystem Research Group (ABERG; andesbiodiversity.org) plot network in Manu National Park, Peru (Figure 1). This site is located on the eastern slope of the tropical Andes in southeastern Peru and is a 3.5 km elevational gradient from the Andean tree line to the Amazonian lowlands. With 23-1 ha and 33-0.1 ha forest plots along a series of descending ridges, this is the most continuous transect in the tropical Andes. To date, ∼19,000 individual trees have been documented in these plots from 100 families, 292 genera, and 1255 species (Farfan-Rios et al. 2025).

Intensive field sampling was conducted for *Prunus* species along this transect, with botanists searching in lateral transects centered on the 1-ha plots (approximately every 250 m) for up to 6 hrs. Leaf tissue was field collected and stored in silica following the Smithsonian Institution guidelines for collecting vouchers and tissues intended for genomic work (Funk et al. 2017). Samples were also included for species known to co-occur across an additional plot network spanning 4 km of elevation in Madidi National Park, Bolivia, and from at least one external population when possible. These samples were collected from herbarium specimens previously not treated with alcohol or fresh tissue stored in silica collected from individuals in the Cotapata region of Bolivia (Figure 1, Table S1). In total, we sampled 158 individuals from 13 species including two non-Andean racemose species (*P. myrtifolia* and *P. caroliniana*) and a reference (Table S1). At least four of these species have not been included in any previous phylogeny.

### ddRAD-seq data

High molecular weight DNA was extracted from ∼50 mg of dried leaf tissue using Aboul-Maaty and Oraby’s (2019) modified CTAB protocol for non-model plants with a few modifications: samples were ground with a bead-beater, incubated at -20 C° overnight (and up to 24 hrs), and the DNA pellet was washed twice with 500 µl 70% ethanol. DNA was checked for quality and quantity on a 1.5% agarose gel and Qubit before being sent to Floragenex (Beaverton, Oregon, USA) for ddRAD library prep and sequencing. Samples were ligated with 6 bp barcodes and digested with the PstI/MseI+0 enzyme pair. Genomic data were demultiplexed and assembled to sample using ipyrad v. 0.9.95 (Eaton and Overcast 2020) with an average of 2,215,444 ± 1,419,356 million reads (mean ± s.d.) recovered per sample.

Reads were assembled into orthologous loci using reference-based assembly in ipyrad. Default settings were used to trim and filter reads and the clustering threshold was set to 0.85 as recommended by the ipyrad manual and McCartney-Melstad et al. (2019) for phylogenetic analyses. For the reference-based assembly, reads were mapped to the *Prunus persica* (peach) genome (GCF_000346465.2; The International Peach Genome Initiative et al. 2013, Verde et al. 2017). The proportion of mapped reads varied across samples (mean = 67%, s.d. = 13%), but variation in the quality of sample preservation and DNA extractions likely explain much of the variation. When calling consensus alleles, we allowed for up to 4 alleles per sample following Donoghue et al. (2022), as all tropical racemose *Prunus* species studied to date are tetraploids (2n=4x=32; Zhao et al. 2016, Hodel et al. 2023, Xie et al. 2025). Cytogenetic data is not available for most species included here, but the clade is thought to derive from an ancient allopolyploid event (Hodel et al. 2023, Xie et al. 2025). Therefore, residual uncertainty in allele dosage and homeologous variation may remain. To evaluate the sensitivity of downstream results to allele representation, we also generated an artificially diploidized dataset by randomly downsampling to a maximum of two alleles per site per sample, but the qualitative patterns were like those represented by the full polyploid-aware assembly. The final reference-mapped dataset comprised 35,099 loci, representing non-overlapping genomic regions where data from at least 4 samples passed filtering. Further downstream analyses were performed using vcftools and the ipyrad-analysis toolkit, which allows for filtering and subsampling loci, genomic windows, and/or SNPs for phylogenetic inference and D-statistics.

### Individual-level phylogeny

A maximum likelihood (ML) phylogeny was reconstructed from a subset of representative samples with the most data present per species (up to 10 individuals per species, at least 1000 loci per sample; n=47), including conspecifics from multiple localities when possible. The ipa.window_extracter module in the ipyrad-analysis toolkit was used to extract data from the 8 largest scaffolds (the chromosomes) and filter sites to only maintain those shared by at least 25% of individuals. We selected the 25% threshold after testing various other filtering thresholds (10%, 30%, 40%, 50%), as 25% provided for the best phylogenetic resolution while also including as much information as possible. The filtered and concatenated sequence matrix is 2.8 Mb in length, containing 107,708 SNPs and 65.8% missing data. This alignment was used in RAxML (Stamatakis 2014) under the GTR+GAMMA evolutionary model to infer a best phylogeny under the rapid hill-climbing algorithm, with support values from 100 bootstrap replicates. The phylogeny was rooted to the reference genome *P. persica* (a native of China and member of the Solitary-flower group and Amygdalus subgenus) and included two additional non-Andean racemose species—*P. caroliniana* (native of the eastern United States and Carribbean) and *P. myrtifolia* (native to the Caribbean, Central America, and lowland South America).

### Species-level phylogeny

To construct the species-level phylogeny, we selected one representative specimen for each species in the individual-level phylogeny (n=13). The samples selected grouped with other members of their species on the individual-level phylogeny and had the most data present. For the species that were reconstructed as non-monophyletic on the individual-level tree, individual selection did result in some movement on the species-level phylogeny. However, we found this was largely linked to amount of missing data and therefore relied on the individual with the highest genomic coverage as the representative. Again, we used the ipa.window_extracter module in the ipyrad-analysis toolkit to extract data from the chromosomes and filter sites, selecting those shared by at least 25% of all species (after testing multiple filtering thresholds). The resulting sequence matrix is 4.2 Mb with 62,262 SNPs and 63.6% missing data. To reconstruct the phylogenetic relationships, we used both maximum likelihood (RAxML) and Bayesian (ExaBayes) analyses. We ran RAxML using GTR+GAMMA to infer a best phylogeny under the rapid hill-climbing algorithm with support values from 100 bootstrap replicates. For ExaBayes, we ran two Metropolis-coupling replicates with four coupled-chains (each with three heated chains) for 1×10^6^ MCMC generations, sampled every 500 generations. Estimated sample size (ESS) for all parameters and branch lengths were summarized with the ProcParam tool and confirmed as sufficiently sampled. The majority-rule consensus phylogeny was generated with the ExaBayes ‘consense’ tool with a 25% burn-in.

To assess the robustness of the resolved relationships and distinguish between conflicted and poorly informed branches, we used Quartet Sampling (Pease et al. 2018; https://www.github.com/ fephyfofum/quartetsampling). We performed Quartet Sampling on the resulting RAxML tree with the full concatenated alignment, allowing 100 replicates per branch (Figure S2).

### Divergence time dating

To assess the timing of divergence among species, we generated a time-calibrated phylogeny using BEAST v. 2.7.7 (Bouckaert et al., 2019) with the same alignment used for the species-level phylogenies. We used an optimized relaxed molecular clock and Yule speciation process with the GTR+GAMMA substitution model. A nodal calibration was set for the crown age of *Prunus* using a normal prior with an offset of 61.5 Mya and a standard deviation of 3.0 Mya (Chin et al. 2014; Su et al. 2023). The crown age of the racemose group was calibrated from a staminate flower of *Prunus hirsutipetala* D.D. Sokoloff, Remizowa et Nuraliev preserved in the late Eocene (Priabonian) Rovno amber from northwestern Ukraine (Sokoloff et al. 2018), using a normal prior with an offset of 35.0 Mya and standard deviation of 2.0 Mya (following Su et al. 2023). To ensure a more rapid convergence of MCMC runs, we used our RAxML topology as our starting tree. The Markov chain Monte Carlo (MCMC) run was conducted for 20,000,000 generations and trees sampled every 2000 generations producing a total of 10,000 trees. The resulting log files were checked in Tracer v.1.5 (Rambaut et al. 2018) and we used TreeAnnotator v.1.10.4 to generate the maximum clade credibility tree discarding 25% of trees as burn-in before visualizing the results in FigTree v.1.4.4 (http://tree.bio.ed.ac.uk/software/figtree/).

### Tests of historical introgression

To test for historical introgression, we used the four-taxon Patterson’s *D*-statistic (ABBA-BABA), which quantifies allele frequency patterns that are incongruent with the hypothesized species tree topology (Green et al. 2010). We ran the Patterson’s *D-*tests using the ipa.baba tool as implemented in ipyrad, which is specifically designed for analysis of RAD-seq datasets. We used the species-level RAxML tree as our starting tree hypothesis and constrained the tests setting the outgroup to *P. myrtifolia*, so all tests would focus on ingroup Andean species. To assess significance, we performed 1000 bootstrap replicates for each test with loci resampled with replacement. Z-scores were then calculated as the number of bootstrap standard deviations from the null *D*=0 (Table 1). Z-scores > 2.5 were considered significant.

**Table 1.** Summary of significant D-statistic tests, with *P. myrtifolia* set to P4.

| P1 | P2 | P3 | dstat | Z | ABBA | BABA | nloci |
| --- | --- | --- | --- | --- | --- | --- | --- |
| *P. antioquensis* | *P. stipulata* | *P. debilis* | -0.69 | 4.75 | 8.38 | 45.13 | 152 |
| *P. stipulata* | *P. antioquensis* | *P. debilis* | 0.69 | 4.60 | 45.13 | 8.38 | 152 |
| *P. amplifolia*, *P. stipulata* | *P. antioquensis* | *P. debilis* | 0.66 | 4.16 |  |  |  |
|  |  |  |  |  | 43.19 | 8.94 | 163 |
| *P. antioquensis* | *P. amplifolia*, *P. stipulata* | *P. debilis* | -0.66 |  |  |  |  |
|  |  |  |  | 4.01 | 8.94 | 43.19 | 163 |
| *P. stipulata* | *P. pleiantha* | *P. detrita*,<br>*P. debilis* | 0.13 | 2.94 | 793 | 605.44 | 3968 |
| *P. pleiantha* | *P. stipulata* | *P. detrita*,<br>*P. debilis* | -0.13 | 2.87 | 605.44 | 793 | 3968 |
| *P. herthae*, *P. huantensis* | *P. pleiantha* | *P. detrita*,<br>*P. debilis* | 0.11 | 2.81 | 864.08 | 686.51 | 4485 |
| *P. stipulata* | *P. huantensis*,<br>*P. antioquensis* | *P. pleiantha* | 0.09 | 2.71 | 1190.20 | 989.74 | 6218 |
| *P. huantensis*,<br>*P. antioquensis* | *P. stipulata* | *P. pleiantha* | -0.09 | 2.71 | 989.74 | 1190.2 | 6218 |
| *P. pleiantha* | *P. herthae*, *P. huantensis* | *P. detrita*,<br>*P. debilis* | -0.11 | 2.67 | 686.51 | 864.08 | 4485 |
| *P. stipulata* | *P. pleiantha* | *P. detrita* | 0.13 | 2.66 | 744.38 | 578.5 | 3725 |
| *P. pleiantha* | *P. antioquensis* | *P. debilis* | 0.51 | 2.63 | 20.25 | 6.5 | 142 |
|  |  |  | -0.13 |  |  |  |  |
| *P. pleiantha* | *P. stipulata* | *P. detrita* |  | 2.62 | 578.5 | 744.38 | 3725 |
| *P. antioquensis* | *P. pleiantha* | *P. debilis* | -0.51 | 2.58 | 6.5 | 20.25 | 142 |
| *P. pleiantha* | *P. stipulata* | *P. debilis* | -0.27 | 2.53 | 68.5 | 119.25 | 576 |
| *P. herthae*, *P. huantensis*, *P. antioquensis*, *P. brittoniana* |  |  | 0.11 |  |  |  |  |
|  | *P. pleiantha* | *P. detrita* |  | 2.52 | 813.45 | 654.78 | 4203 |

### Tests of recent admixture

To assess genetic structure and test for recent admixture, we used the Bayesian clustering algorithm STRUCTURE v. 2.3.4 (Pritchard et al. 2000) as implemented in ipyrad with the set of samples included in the individual-level phylogeny (minus the reference and non-Andean species). SNPs were filtered conservatively to only include those with at least 50% coverage across individuals and then randomly sampled to one SNP per locus to avoid linkage. This resulted in a total of 1493 SNPs included in the analysis. The model was run with a burn-in of 125,000 and run length of 500,000 generations, testing K=1–10 with 10 replicates per K. The assignment of samples into K distinct genetic groups was assessed using DeltaK, which has been shown to be more robust to accurately estimating the correct number of clusters in STRUCTURE, particularly with high amounts of missing data as is common in RADseq (Evanno et al. 2005; Figure S3).

### Estimates of genomic diversity

Genetic diversity was estimated within individuals and species to compare to the admixture results. In VCFtools v.1.16 (Danecek et al. 2011), sequences were filtered using two different missing data thresholds, retaining loci present within at least 50% or 70% of the samples for each species. Sites were also filtered if they had a minor allele frequency less than 0.05. We calculated nucleotide diversity (*π*) for each species – the average pairwise nucleotide differences between sequences in a sample at a site as well as observed (*H_o_*) and expected heterozygosity (*H_e_*) – the proportion of loci where an individual is heterozygous vs. the proportion expected by Hardy-Weinberg equilibrium – within samples and averaged across the species. For these calculations, *P. debilis* and *P. detrita* samples were grouped together as well as *P. huantensis* and *P. brittoniana,* given that STRUCTURE (Figure 4) did not detect any distinct genetic clusters among these species’ pairs. Species with <3 samples were excluded from these analyses given the lack of statistical power from such a small sample size.

## RESULTS

### Phylogenomic relationships and divergence time estimation

Both the species-level maximum likelihood (RAxML) and Bayesian (ExaBayes) phylogenetic analyses supported the same evolutionary relationships among the focal Andean species and the non-Andean lineages (Figure 2). Subtropical *P. caroliniana* was reconstructed as sister to the tropical species and *P. myrtifolia* was sister to the Andean species. Among the partially sympatric Andean species, the lineage including pre-montane species *P. debilis* and *P. detrita* were sister to the remaining Andean species, followed by the mid-elevation cloud-forest species (*P. herthae, P. pleiantha, P. stipulata, P. integrifolia*) with the high-elevation species being the most recently derived (*P. huantensis, P. brittoniana, and P. antioquensis*). Statistical support for the inferred relationships was assessed using bootstrap support (RAxML), posterior probabilities (ExaBayes), and Quartet Sampling.

**Figure 2.**
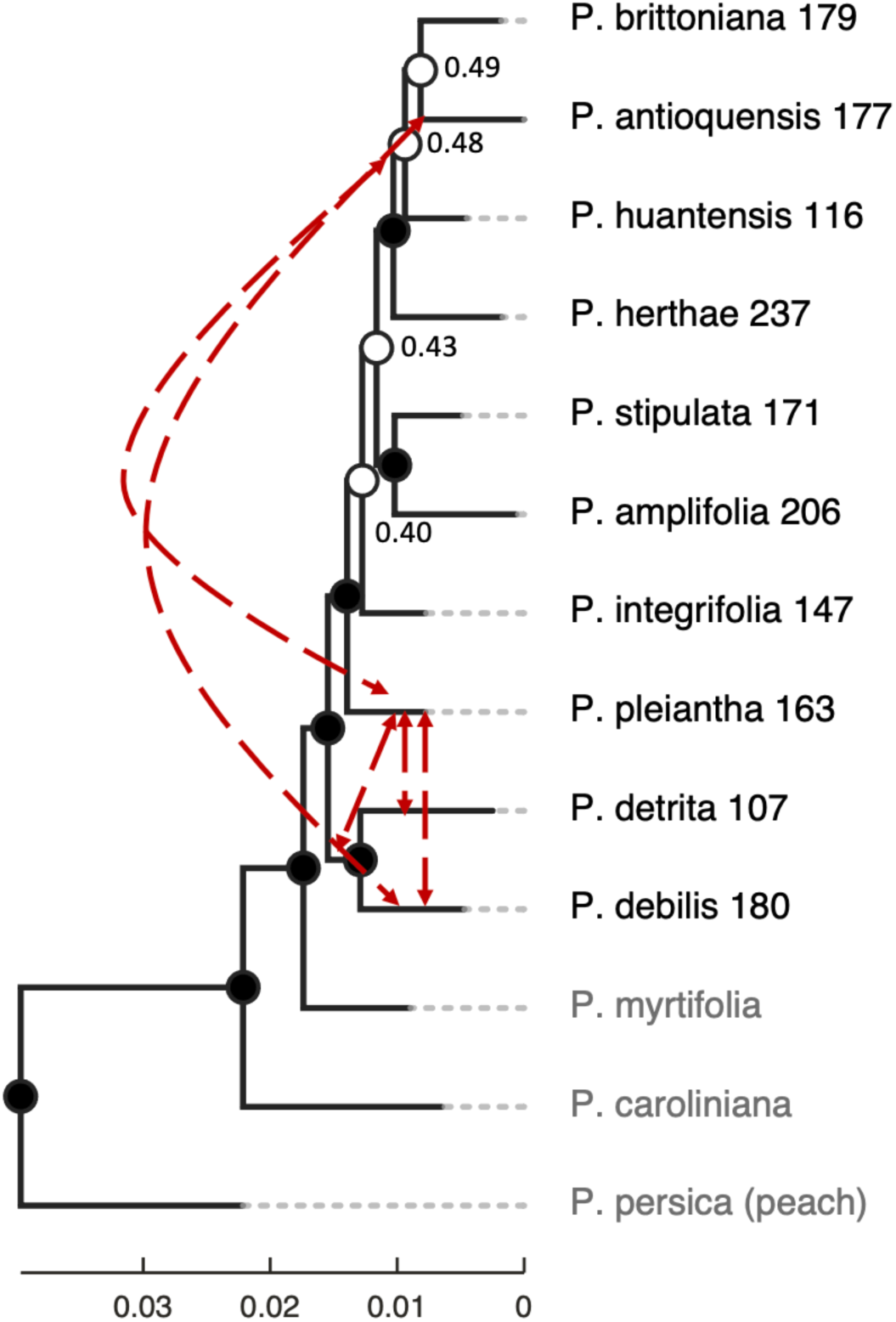
Species-level topology resulting from RAxML and ExaBayes phylogenetic analyses. Nodes show RAxML bootstrap support values where black circles indicate values > 75 and values < 75 are shown on white nodes. All Bayesian posterior probabilities were equal to 1. The tree is rooted to the reference (*P. persica*) and all three non-Andean species are shown in grey. Dashed arrows summarize patterns of historical admixture inferred from D-statistics.

Quartet Sampling (QS; Pease et al. 2018) complements conventional node-support metrics by distinguishing low support caused by limited phylogenetic information from low support caused by conflicting signal. It evaluates each node using the relative frequency of alternative relationships among sets of four taxa. In this framework, QI (quartet informativeness) measures the proportion of sampled quartets that are informative for a given node, QC (quartet concordance) measures the degree to which informative quartets support the focal relationship, and QD (quartet differential) measures the skew between the two alternative discordant topologies, helping distinguish balanced discordance (as expected under incomplete lineage sorting) from a strong preference for one alternative (consistent with introgression). While the tree nodes are generally well-supported (all posterior probabilities = 1, most bootstrap values >75; Figure 2), Quartet Sampling indicated that low bootstrap support values are not likely the result of insufficient phylogenetic signal, as QI ranged from 80–100%. The lack of a strong skew in discordant quartets at most nodes (QD>0.5) suggests that incomplete lineage sorting (ILS) is more likely than introgression. In contrast, the branch subtending *P. huantensis, P. brittoniana*, and *P. antioquensis* showed moderate concordance (QC=0.21) and strong discordant skew (QD=0.22), signal consistent with a possible introgressive event (Figure S2).

The time-calibrated phylogeny generated in BEAST recovered the same tree topology as RAxML and ExaBayes with similar branch support (Figure 3). The estimated divergence time *for P. myrtifolia* and the ancestor of the Amazonian and Andean species is ∼27.3 Ma (95% highest posterior density [HPD] 22.4–31.7). The Andean species began diverging at the start of the Miocene, with the pre-montane *P. debilis*+*detrita* lineage branching off ∼23.5 Ma (95% HPD 19.4-28.1), and from each other ∼17.6 Ma (95% HPD 12.8-22.6). The cloud forest species followed, with the *P. pleiantha* lineage diverging ∼20.4 Ma (95% HPD 16.3-24.4), and *P*. *integrifolia* ∼18.0 Ma (95% HPD 14.2-21.7). The remaining lineages emerged in the mid to late-Miocene (15.9–9.3) with *P. huantensis* ∼11.6 Ma (95% HPD 8.6-14.9) and *P. antioquensis*+ *brittoniana* ∼9.3 Ma (95% HPD 6.2-12.3). Together, the phylogenetic analyses indicate that these lineages are ancient, with potential sympatry since the Miocene.

**Figure 3.**
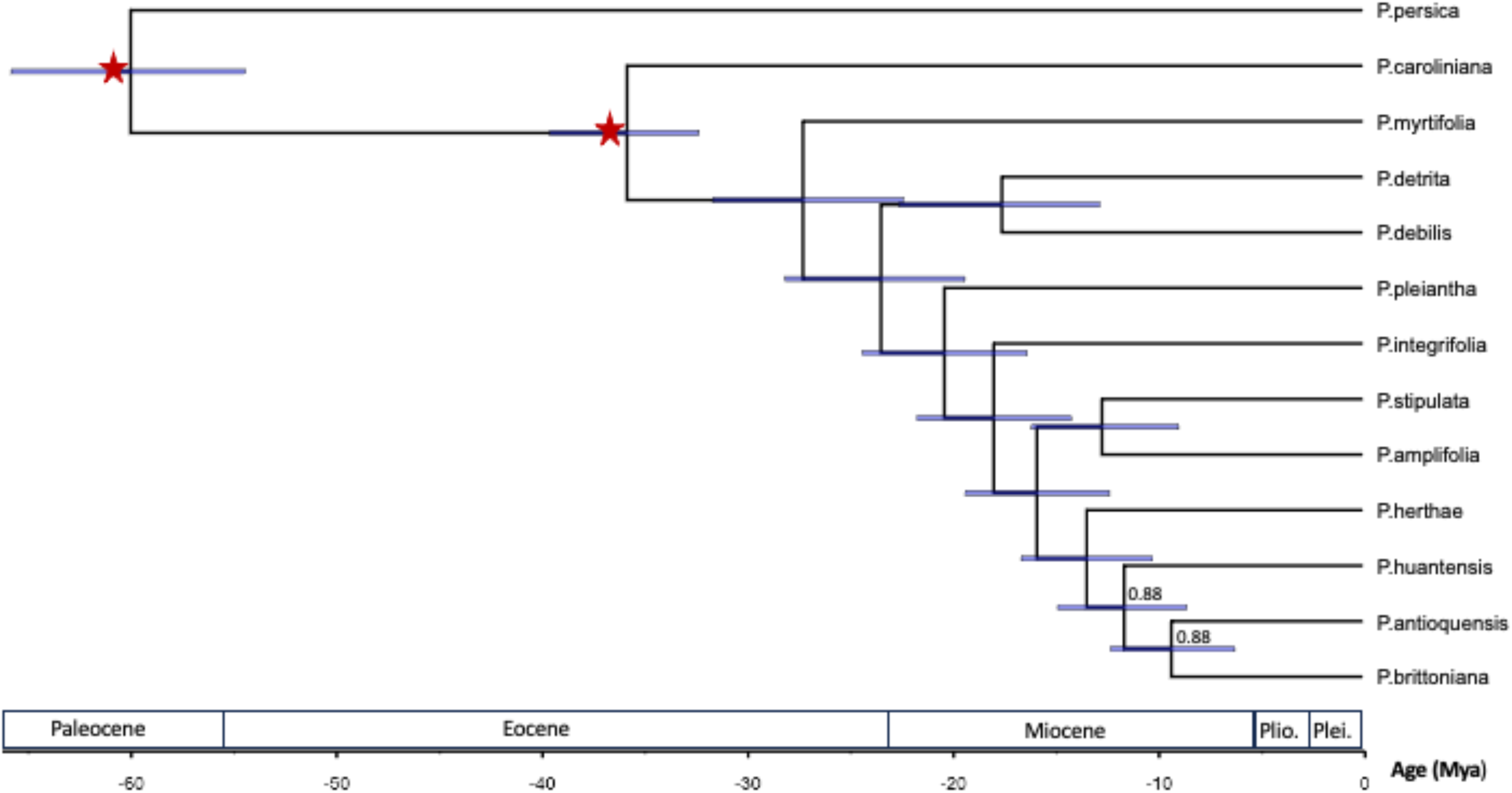
Time-calibrated phylogeny generated from BEAST analysis. Posterior probabilities are shown for nodes when < 1. Node bars represent the 95% highest probability density (HPD) interval for node ages. Red stars indicate the fossil calibration points.

**Figure 4.**
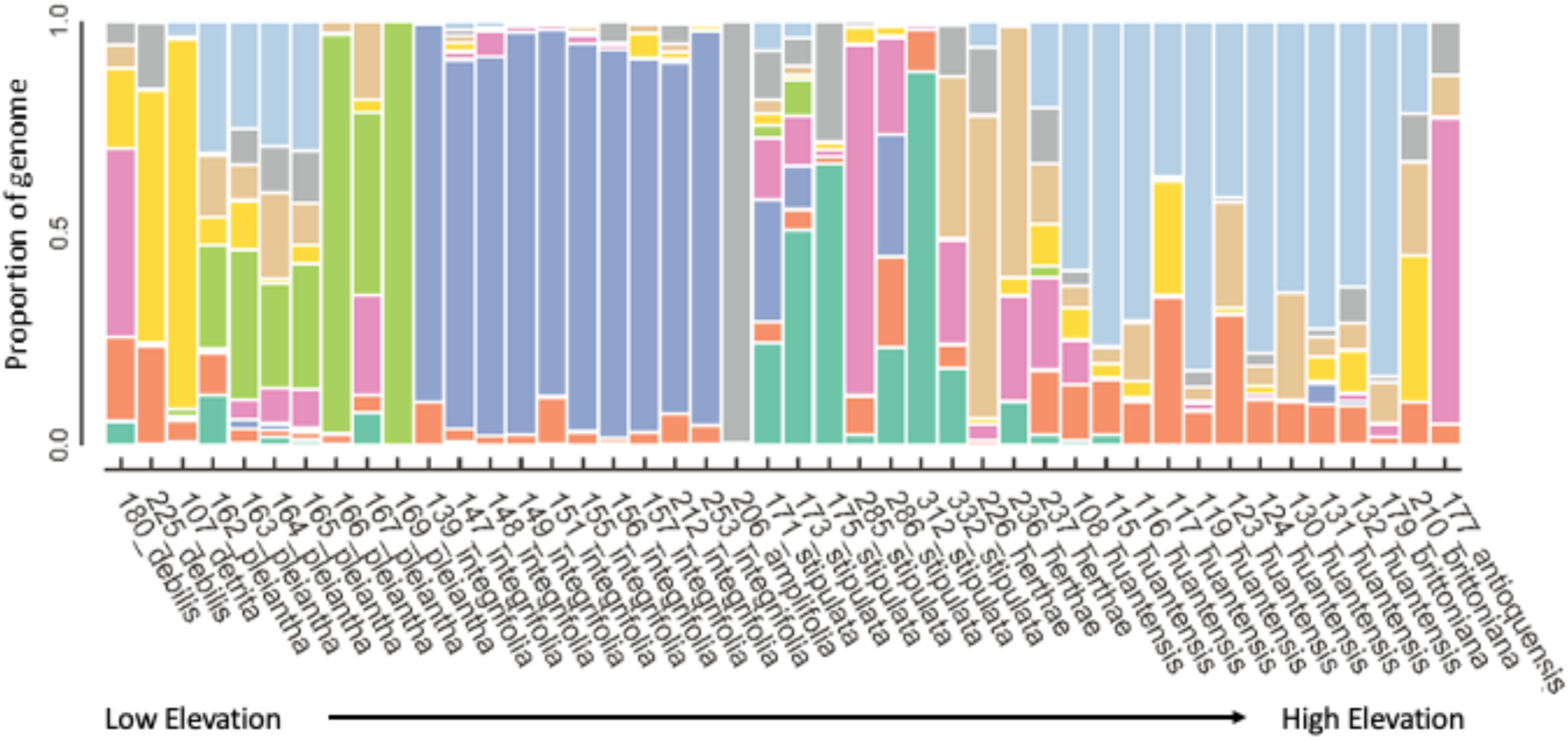
Patterns of genetic structure among individuals of sympatric species. Each vertical bar represents a single individual and the y-axis indicates the proportion of genetic ancestry of that individual assigned to different inferred genetic clusters, designated with different colors. Species are ordered on the x-axis according to their elevational range, from low to high. STRUCTURE detected K=9 distinct genetic clusters as the most likely grouping (Figure S3).

### Tests of historical introgression and recent admixture

*D*-statistics indicated significant historical introgression events between *P. pleiantha* and the ancestor of *P. debilis+detrita*, with more limited evidence suggesting introgression with *P. debilis* and *P. detrita* separately (Figure 2, Table 1). A second significant introgression event was detected between *P. pleiantha* and the *P. huantensis*+*antioquensis* ancestor, which is further supported by Quartet Sampling (Figure S2). Finally, significant *D*-statistics supporting introgression between *P. debilis* and *P. antioquensis* were also detected. Overall, introgression is widespread and does not seem to be limited by ecological or evolutionary distance, as evidenced by their elevational niche overlap (Figure 1) and hypothesized phylogenomic relationships (Figure 2).

STRUCTURE analysis found K=9 to be the most likely number of distinct genetic clusters within the dataset, determined by mean log probability and DeltaK values (Figures 5, S3). Individuals largely clustered according to their morphologically assigned species; however, the number of detected clusters was fewer than the expected number of named taxonomic species in the analysis (*n*=10). *P. integrifolia* and *P. amplifolia* are largely reproductively isolated, though low levels of admixture were detected in *P. integrifolia* and only one sample was included for *P. amplifolia.* Admixture is widespread amongst all other species, particularly *P. pleiantha*, *P. stipulata*, *P. herthae*, *P. debilis, P. huantensis, P. brittoniana and P. antioquensis,* and like the introgression analyses, appears unrelated to ecological or evolutionary distance.

### Estimates of genomic diversity

Within each species, the individuals with the highest observed heterozygosity (*H_o_*) were also the most admixed in the STRUCTURE results (Tables 3 and 4, Figure 4), and this was consistent with different amounts of missing data. For example, in *P. pleiantha*, individuals 163, 164, and 165 showed high levels of admixture from multiple groups and had the highest *H_o_* estimates (50%: 0.18 – 0.22, 70%: 0.19 – 0.25; Table 3). Across species and filtering thresholds, mean pairwise nucleotide diversity (*π*) was high (50%: 0.41 – 0.64, 70%: 0.37 – 0.51) compared to *H_o_*, *π* tracked with *H_e_* as expected, and *H_o_* (50%: 0.06 – 0.15, 70%: 0.06 – 0.17) was far below *H_e_* (50%: 0.41 – 0.64; 70%: 0.37 – 0.51; Table 2). However, the estimates for *P. herthae* and the *P. debilis*+*detrita* group should be interpreted cautiously, given the low sample sizes for each. These statistics corroborate the findings of the STRUCTURE analysis and further indicate a history of interspecific gene flow among co-occurring *Prunus* species.

**Table 2.** Summary of within species genomic diversity statistics.

| Species | # of samples | Filter | # of loci | Average $\pi$ | Average $H_o$ | Average $H_e$ |
| --- | --- | --- | --- | --- | --- | --- |
| *P. integrifolia* | 10 | 50 | 53574 | 0.42 ± 0.13 | 0.15 ± 0.064 | 0.42 ± 0.0025 |
| | | 70 | 17592 | 0.41 ± 0.14 | 0.17 ± 0.073 | 0.40 ± 0.0015 |
| *P. huantensis*/<br>*brittoniana* | 12 | 50 | 24457 | 0.41 ± 0.13 | 0.12 ± 0.078 | 0.41 ± 0.0012 |
| | | 70 | 1960 | 0.38 ± 0.15 | 0.16 ± 0.10 | 0.37 ± 0.0031 |
| *P. stipulata* | 7 | 50 | 24457 | 0.45 ± 0.11 | 0.095 ± 0.096 | 0.45 ± 0.0024 |
| | | | | 0.42 ± 0.13 | 0.11 ± 0.11 | 0.42 ± 0.0064 |
| *P. pleiantha* | 7 | 50 | 23299 | 0.45 ± 0.11 | 0.12 ± 0.081 | 0.44 ± 0.0015 |
| | | 70 | 7295 | 0.43 ± 0.12 | 0.13 ± 0.091 | 0.42 ± 0.0013 |
| *P. herthae* | 3 | 50 | 6251 | 0.62 ± 0.077 | 0.085 ± 0.03 | 0.61 ± 0.0067 |
| | | 70 | 699 | 0.51 ± 0.073 | 0.064 ± 0.0072 | 0.51 ± 0 |
| *P. debilis*/*detrita* | 3 | 50 | 1747 | 0.64 ± 0.061 | 0.064 ± 0.058 | 0.064 ± 0.0009 |

**Table 3.** Summary of within-individual heterozygosity estimates.

| Species | Individual | H<sub>o</sub> (50% filter) | H<sub>e</sub> (50% filter) | H<sub>o</sub> (70% filter) | H<sub>e</sub> (70% filter) |
| --- | --- | --- | --- | --- | --- |
| *P. integrifolia* | 139_integrifolia | 0.11 | 0.42 | 0.12 | 0.41 |
|  | 147_integrifolia | 0.31 | 0.42 | 0.36 | 0.4 |
|  | 148_integrifolia | 0.14 | 0.42 | 0.16 | 0.4 |
|  | 149_integrifolia | 0.18 | 0.42 | 0.21 | 0.4 |
|  | 151_integrifolia | 0.10 | 0.42 | 0.12 | 0.4 |
|  | 155_integrifolia | 0.16 | 0.42 | 0.19 | 0.4 |
|  | 156_integrifolia | 0.12 | 0.42 | 0.14 | 0.4 |
|  | 157_integrifolia | 0.17 | 0.42 | 0.2 | 0.4 |
|  | 212_integrifolia | 0.09 | 0.41 | 0.1 | 0.4 |
|  | 253_integrifolia | 0.12 | 0.42 | 0.14 | 0.41 |
| *P. huantensis*<br>*/brittoniana* | 108_huantensis | 0.16 | 0.41 | 0.23 | 0.37 |
|  | 115_huantensis | 0.1 | 0.41 | 0.13 | 0.38 |
|  | 116_huantensis | 0.22 | 0.41 | 0.29 | 0.37 |
|  | 117_huantensis | 0.044 | 0.4 | 0.07 | 0.37 |
|  | 119_huantensis | 0.072 | 0.41 | 0.093 | 0.37 |
|  | 123_huantensis | 0.069 | 0.4 | 0.12 | 0.37 |
|  | 124_huantensis | 0.2 | 0.4 | 0.25 | 0.37 |
|  | 130_huantensis | 0.05 | 0.41 | 0.079 | 0.38 |
|  | 131_huantensis | 0.15 | 0.41 | 0.22 | 0.37 |
|  | 132_huantensis | 0.26 | 0.41 | 0.36 | 0.37 |
|  | 179_brittoniana | 0.064 | 0.41 | 0.08 | 0.37 |
|  | 210_brittoniana | 0.03 | 0.4 | 0.05 | 0.38 |
| *P. stipulata* | 171_stipulata | 0.28 | 0.45 | 0.32 | 0.42 |
|  | 173_stipulata | 0.17 | 0.45 | 0.18 | 0.42 |
|  | 175_stipulata | 0.07 | 0.45 | 0.07 | 0.41 |
|  | 285_stipulata | 0.058 | 0.45 | 0.1 | 0.42 |
|  | 286_stipulata | 0.03 | 0.45 | 0.041 | 0.41 |
|  | 312_stipulata | 0.037 | 0.45 | 0.032 | 0.42 |
|  | 332_stipulata | 0.02 | 0.45 | 0.021 | 0.43 |
| *P. pleiantha* | 162_pleiantha | 0.12 | 0.44 | 0.13 | 0.43 |
|  | 163_pleiantha | 0.22 | 0.44 | 0.25 | 0.43 |
|  | 164_pleiantha | 0.18 | 0.44 | 0.19 | 0.42 |
|  | 165_pleiantha | 0.2 | 0.44 | 0.22 | 0.42 |
|  | 166_pleiantha | 0.023 | 0.44 | 0.021 | 0.42 |
|  | 167_pleiantha | 0.05 | 0.44 | 0.052 | 0.42 |
|  | 169_pleiantha | 0.051 | 0.44 | 0.056 | 0.42 |
| *P. herthae* | 226_herthae | 0.087 | 0.62 | 0.057 | 0.51 |
|  | 236_herthae | 0.054 | 0.61 | 0.063 | 0.51 |
|  | 237_herthae | 0.11 | 0.62 | 0.072 | 0.51 |
| *P. debilis*/<br>*detrita* | 180_debilis | 0.13 | 0.64 | NA | NA |
|  | 225_debilis | 0.036 | 0.64 | NA | NA |
|  | 107_detrita | 0.026 | 0.64 | NA | NA |

## DISCUSSION

In this study, we aimed to characterize the evolutionary relationships among congeneric tree species that turn over with elevation in the tropical Andes, assess whether they are genetically isolated, and the extent to which gene flow is related to ecological or evolutionary distance. The phylogenomic data presented here suggest that *Prunus* species known to co-occur as elevational replacements diversified across the Miocene (∼23.5-9.3 Ma). However, despite these lineages being separated by considerable evolutionary time, species are not reproductively isolated, as substantial gene flow is detectable among species through time and does not decrease with increasing evolutionary distance or elevational separation. Together, these findings suggest that during diversification, observable ecological and morphological differences may emerge despite ongoing gene flow. Further, the maintenance of incomplete reproductive barriers over millions of years may be an important and overlooked mechanism for maintaining high tree diversity, particularly with the low population densities and high environmental variability characteristic of tropical montane forests.

### Gene flow through time

Our genomic analyses show that Andean *Prunus* species remain distinct but porous evolutionary lineages. Despite deep Miocene divergences, we detect both ancient introgression and recent admixture among partially sympatric taxa. *D*-statistics detected at least five significant ancient introgression events. Two of these events occurred among lineages separated by great evolutionary and elevational distance (*P. pleiantha* × *P. huantensis+ antioquensis*, and *P. debilis* × *P. antioquensis;* Figure 2). Introgression was also detected between *P. pleiantha* and the ancestor of *P. debilis+ detrita* (*P. pleiantha* × *P. debilis+ detrita*) as well as among *P. pleiantha* × *P. debilis* and *P. pleiantha* × *P. detrita* individually. These patterns are most consistent with repeated interspecific introgression among the ancestors of these lineages, with gene flow continuing as elevationally separated lineages diverged. However, because D-statistics do not resolve the direction of introgression, a stronger test of this interpretation will require methods that can distinguish donor and recipient lineages (such as D-FOIL; Pease and Hahn 2015), together with denser taxonomic and wider genomic sampling.

Historical introgression among species separated in ecological space across the elevation gradient is plausible for several reasons. While the medians of their elevation ranges may be quite different, they are found in close geographic proximity, and their elevational ranges overlap (or are much closer) at the range margins. *Prunus* species are thought to be obligate outcrossers with generalist pollinators, as is common in tropical trees (Ashton 1969, Bawa 1974, 1979). The extent of self-incompatibility across the genus and potential for breakdown after polyploidization are still unclear (Hauck et al. 2006, Tao et al. 2010, Aguiar et al. 2015). Generalist pollinators can forage over kilometers, providing opportunity for gene flow, especially given that many tropical species flower semi-synchronously during the dry season (Bawa et al. 1985, Van Schaik et al. 1993).

Even where known species ranges are separate at present, they have not necessarily always been so. Pollen records from Lago Consuelo (1360 m a.s.l., located in southeastern Peru adjacent to the Bolivian border) show that during the last full glacial period (43.5–22 kyr BP), individual cloud forest species migrated downslope at least 1000–1700 vertical m in response to lower cloud base and cooler temperatures (-5°C; Urrego et al. 2005). Similar pulses of connectivity likely occurred as many as ten times in response to glacial-interglacial cycles over just the last million years and may have helped maintain these incomplete reproductive boundaries. Today, these species ranges are separated as far apart as they have likely been during the Quaternary, and historical introgression events may be signatures of past ecological overlap.

Tests of recent admixture also found gene flow to be widespread among Andean *Prunus* species. *P. integrifolia* and *P. amplifolia* are largely genomically isolated, but gene flow appears rampant among all other species. The isolation of *P. integrifolia* is particularly interesting as this species has the widest elevational range and is one of the most abundant species in the tropical Andes. The single sample of *P. amplifolia* included in the analysis prevents us from determining whether this pattern of isolation is widespread in the species (as observed in *P. integrifolia*) or whether rates of mixed genetic ancestry are highly variable (as observed in *P. pleiantha*). Gene flow was especially high among *P. pleiantha, P. stipulata*, and *P. herthae*, all cloud forest species with largely overlapping elevation ranges. However, individuals of these species mostly form monophyletic groups, indicating they have developed specific genetic distinctiveness as well as morphological similarity that allows for field identification despite gene flow (Figure S1). As *P. pleiantha* was identified in most of the historical introgression events and had high proportions of mixed genetic ancestry, we can infer that this species readily hybridizes when in contact with close relatives. Yet, previous morphological studies of herbarium specimens (Achá 2013, Pérez-Zabala 2022) have not recovered high morphological variation in *P. pleiantha* or *P. herthae*, as would be expected if morphological variation increased with gene flow. Further investigation will be needed to test these hypotheses, identify the directionality of such gene flow, and determine whether physiological, ecological, or genetic factors drive this weaker reproductive isolation and its implications.

These inferences should also be interpreted considering the likely shared ancient allopolyploid origin of the clade. Residual homeologous variation or differential sorting of ancestral subgenomes, may increase apparent allele sharing among taxa and complicate the interpretation of admixture and D-statistic results, particularly in analyses that reduce polyploid variation to diploid-like summaries. However, because this polyploid background is likely shared across the clade, any resulting bias should be widespread rather than specific to particular taxa. Polyploid history may therefore contribute to overall ambiguity in the genomic signal, but is unlikely to fully explain the repeated, lineage-specific patterns of excess allele sharing recovered here.

STRUCTURE did not identify *P. debilis* and *P. detrita* as distinct genetic clusters, possibly due to how closely related these species are, gene flow, and the low number of individuals for both species in the dataset. Our genomic data also support the most recent taxonomic revision by Pérez-Zabala (2022) which considered *P. brittoniana* and *P. huantensis* to be the same species given their morphological and ecological similarities (Figure S1). Taken together, these results indicate that gene flow has been a persistent and complex feature of Andean *Prunus* diversification, occurring across both evolutionary time and ecological space, despite ancient divergence times and emergence of species-level distinctiveness.

### Diversification patterns and biogeographic implications

Phylogenomic reconstructions and divergence time dating indicate that *Prunus* diversified in the tropical Andes over the Miocene. In agreement with prior studies, we found *P. myrtifolia* (endemic to the Caribbean, Central America, and lowland South America) to be sister to the Andean species included in this analysis, supporting the hypothesis that this lineage or its ancestor colonized South America via migration through the Caribbean and Central America (Chin et al. 2014, Hodel et al. 2023). While our divergence time estimate for *P. myrtifolia* of ∼27 Ma is slightly later than the Neotropical colonization of ∼34 Ma predicted by Chin et al. (2014), it is still within the climatic shift from warmhouse to transitional state that coincided with an onset of glaciation in Antarctica (∼33.9 – 27.29 Ma; Judd et al. 2024) and the estimated end of the boreotropical forests in the Northern Hemisphere.

Among the Andean species included in our analyses, the two lowest elevation species (*P. debilis* and *P. detrita*) were sister to the rest of the group (diverging ∼23.5 Ma) and the high-elevation clade was the most recently derived (∼11.6 Ma—*P. huantensis*, ∼9.3 Ma—*P. antioquensis* and *P. brittoniana*). This is consistent with the timing of major uplift events and estimated paleo-elevations in the North-Central and Central Andes (Boschman 2021). It also supports the findings of Hodel et al. (2023), which detected a long period of stasis followed by rapid diversification of Neotropical *Prunus* during the Miocene. Although the exact timing, pace, and sequence of Andean orogeny is a subject of ongoing study and is spatially heterogeneous, the late Oligocene and Miocene (∼25-5 Ma) are known to have been a period of intense and accelerated uplift in the Eastern Cordillera of the North-Central and Central Andes (Boschman et al. 2021, Pérez-Escobar et al. 2022).

The lower montane cloud forest species in our analysis (median elevation ranges 1200–2500 m; *P. pleiantha, P. integrifolia, P. herthae, P. stipulata*) diverged during the early to mid-Miocene. This aligns with the predicted origination of cloud forests (Sempere et al. 2005) and the estimated divergence times for other cloud forest lineages (Luebert and Weigend 2014). Andean cloud forests are distinct habitats characterized by persistent cloud cover and immersion that occurs from ∼1200 m to the upper forest limits at ∼3200–3800 m (depending on latitude). Although they are the most speciose habitats in the Andes, persistence in these environments often requires unique adaptations to cope with the reduced light availability, high precipitation, and humidity such as thick, pachyphyllic leaves, low growth rates, and ability to withstand heavy accumulations of epiphytes (Grubb 1977, Rapp and Silman 2012, Fahey et al. 2016, Pérez-Escobar 2022). The highest elevation species in our study (median elevation ranges ≥3000 m; *P. huantensis/ P. brittoniana, P. antioquensis*) diverged during the late Miocene, as the final uplift phase began in the North-Central and Central Andes (Boschman et al. 2021). Species in this elevation zone are exposed to unique high-elevation stressors such as night frost, with temperatures dipping below 0°C, requiring further physiological specialization (Pérez-Escobar 2022).

Given the phylogenomic reconstructions and divergence time estimates, two hypotheses for understanding diversification among Andean *Prunus* emerge. First, these species may be the result of adaptive diversification following Andean uplift. Similar diversifications have been documented in other Andean plant clades in response to novel mountain habitat opportunities (e.g. Melastomataceae—Dudley 1978, wild tomato—Pease et al. 2016). Dating analysis from this and previous studies (e.g. Chin et al. 2014, Hodel et al. 2023) suggest *Prunus* arrived in South America during the early to middle Oligocene (∼34-27 Mya), long before Miocene mountain building brought the central Andes to their modern elevations. However, an alternate and often dominant pattern of diversification in the Andes is one in which extratropical genera emigrate in and colonize elevation bands for which they were pre-adapted (Griffiths et al. 2020, Segovia et al. 2020, Linan et al. 2021). These elevation bands then become fragmented by topography, climatic oscillations, and geological processes, promoting allopatric speciation.

Examples of such latitudinal allopatric speciation have been documented across plant and bird genera (e.g. *Ceroxylon*—Sanín et al. 2016, *Gunnera*—Bacon et al. 2018, *Scytalopus*—Cadena et al. 2020).

Importantly, these modes of speciation (adaptive vs. allopatric) are not mutually exclusive. Post-colonization, diversification may occur both in response to novel habitat opportunities (across elevation) and following isolation events (across latitude), with both mechanisms responsible for generating the immense rates of endemism and diversity characteristic of tropical Andes (e.g. lupines—Hughes and Eastwood 2006, *Hypericum*—Nϋrk et al. 2013, bellflowers—Lagomarsino et al. 2016, orchids—Pérez-Escobar et al. 2017). Disentangling when and how these mechanisms gave rise to the >100 species of South American *Prunus* (Pérez-Zabala 2022) will require denser species sampling than available in this study as well as higher-resolution species distribution data. Within those limitations, our results are most consistent with diversification in Andean *Prunus* being driven by the joint effects of adaptive diversification in response to uplift-driven ecological opportunity and the immigration, sorting, and isolation of lineages adapted to different elevational environments.

### Conclusions

Here, we found evidence of historical introgression, possibly in response to Quaternary range shifts, and widespread admixture among co-occurring *Prunus* species. Yet despite gene flow, these species have maintained distinct evolutionary lineages with identifiable ecological and morphological differences since the Miocene. These results align with the syngameon model, wherein species maintain incomplete reproductive barriers but primarily experience divergent selection (Lotsy 1925, Grant 1971, Cannon and Lerdau 2015, Cervantes et al. 2025, Schley et al. 2025). Though oaks are the classic example (e.g. Cannon and Petit 2020), such genomic mutualism may be an important yet under-appreciated mechanism in the maintenance of diversity in species-rich tropical forests, where individuals predominantly outcross but retain some capacity for self-fertilization and interspecific hybridization at low population densities (Wu 2001, Cannon and Lerdau 2015). This has important implications in the context of climate change, as species are now shifting their ranges in response to changes in temperature and moisture (Feeley et al. 2011, Fadrique et al. 2018). Though most tree species will not be able to migrate fast enough to track climatic shifts and are instead experiencing range retractions, any upslope range shift may increasingly bring congeneric species into contact, especially at sparsely populated range margins where low conspecific density could elevate the likelihood of heterospecific mating and gene flow (Duque et al 2015, Cannon and Lerdau 2015, Farfan-Rios et al. 2025, Quinlan et al. 2025). Further work is required to understand the prevalence of syngameons in tropical forests and their role in species’ responses to rapid environmental change, namely resilience and avoidance of local extinction. Incorporating these complex evolutionary dynamics into demographic models is essential for accurately predicting the futures of Andean forests and effective conservation and management under current climate change.

## ACKNOWLEDGEMENTS

We thank Paul Fine, Michael Anderson, and Regina Joice Cordy for their helpful comments on study design and earlier versions of this manuscript. We are also grateful to Herbario Vargas (CUZ) and Herbario Nacional de Bolivia (LPB) for their assistance with coordination, field work, and access to herbarium material, as well as Christopher Dick and Fairchild Botanical Garden for their donation of samples to this project. Additional thanks to the Andes Biodiversity Ecosystem Research Group (ABERG), and the many students and field technicians involved with this project (especially Andrea Palomino Cardenas, Jesus Manuel Bañon Sanchez, Daire Luciana Lafuente Palie, Paukar Alejandro Vedia Villca, and Felipe Aramburo Jaramillo). This work was supported by NSF LTREB (1754647) and Garden Club of America’s Tropical Botany Fellowship. The material used in this study was collected under the research permits No 18-2019-SERNANP-JEF, No 14-2022-SERNANP-JEF, No 527-2019-MINAGRI-SERFOR-DGGSPFFS, and No D000170-2023-MIDAGRI-SERFOR-DGGSPFFS-DGSPF granted by SERNANP and SERFOR.

## AUTHOR CONTRIBUTIONS

EJQ, MRS, and JBP initially conceived of the study. EJQ acquired funding, led research design, data collection, data analysis and interpretation, and wrote the manuscript. JSB, AFF, and WFR assisted with data collection and WFR additionally contributed to manuscript editing and translated the summary. JBP assisted with data analysis and manuscript review and editing. MRS provided supervision, acquired funding, and contributed to research design and manuscript writing.

## DATA AVAILABILITY

The genomic data generated for this study are available on NCBI’s GenBank under submission ID SUB15831409 and corresponding herbarium vouchers are deposited in Herbario Vargas (CUZ) at the Universidad Nacional de San Antonio Abad del Cusco (UNSAAC) and Herbario Nacional de Bolivia (LPB).

## SUPPORTING INFORMATION

**Table S1.**
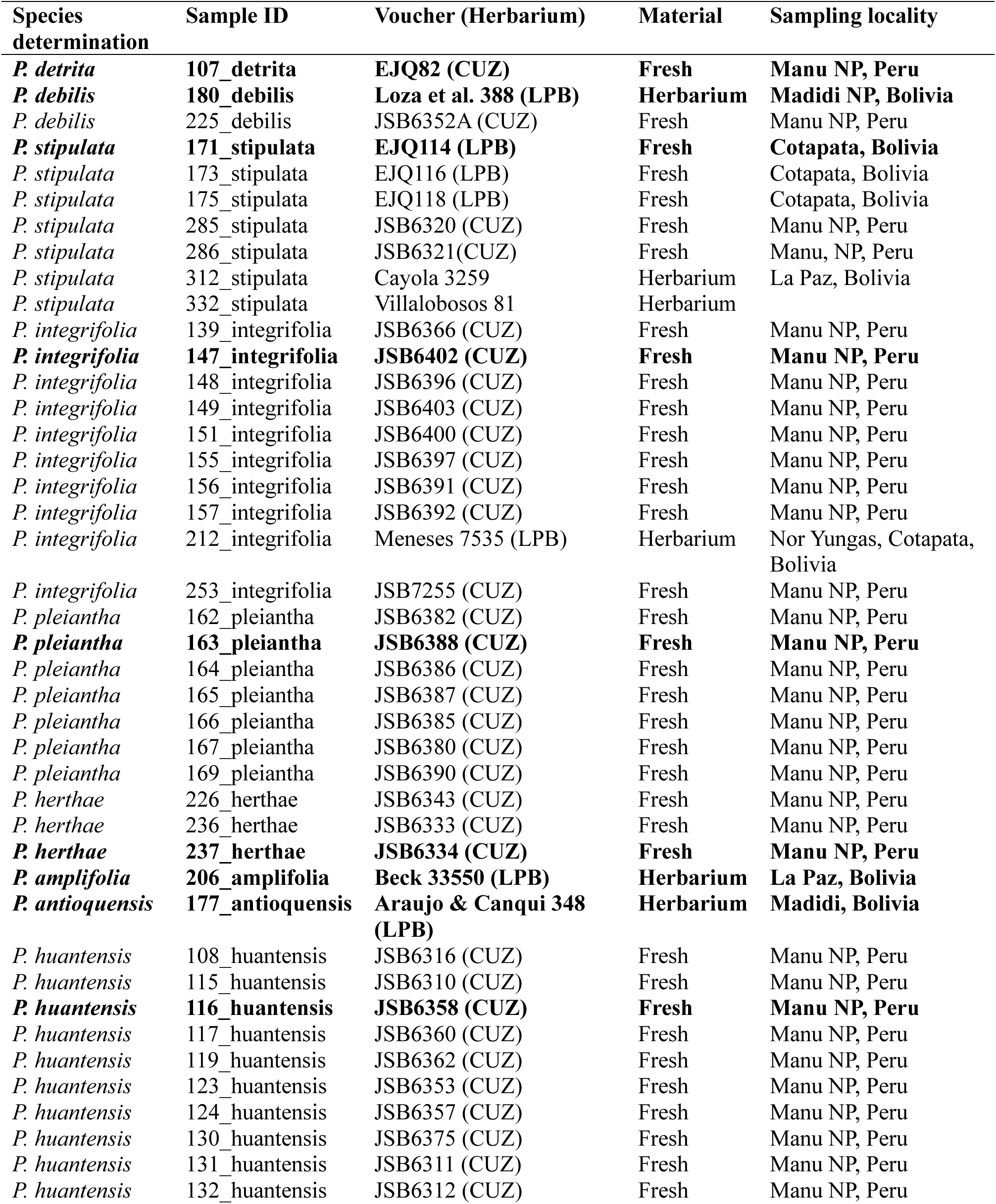
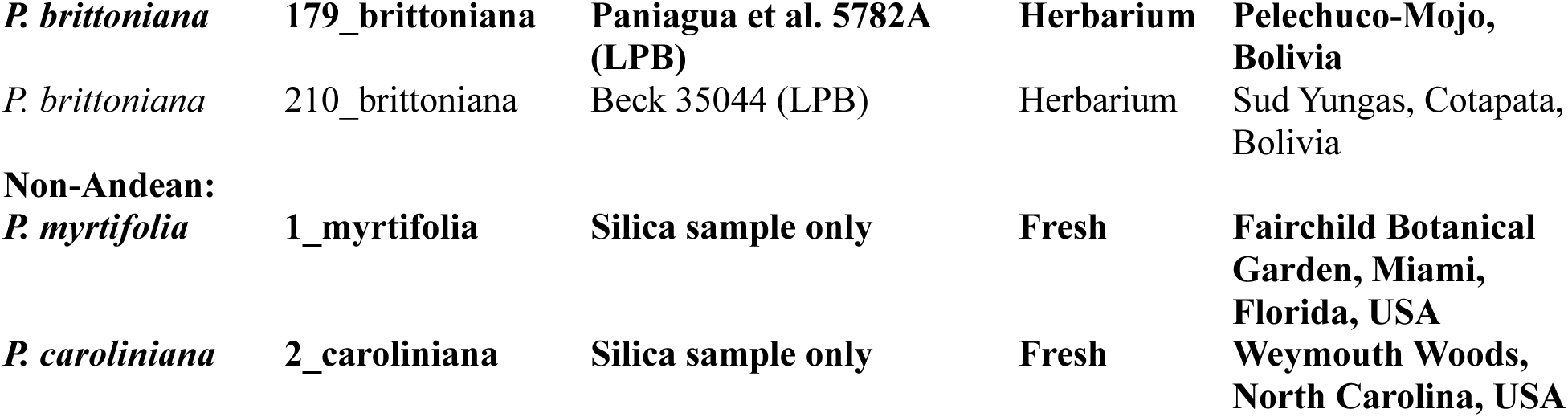
Summary of samples included in individual-level phylogeny and genetic structure analyses. Samples in bold were used to construct the species-level phylogeny.

**Figure S1.**
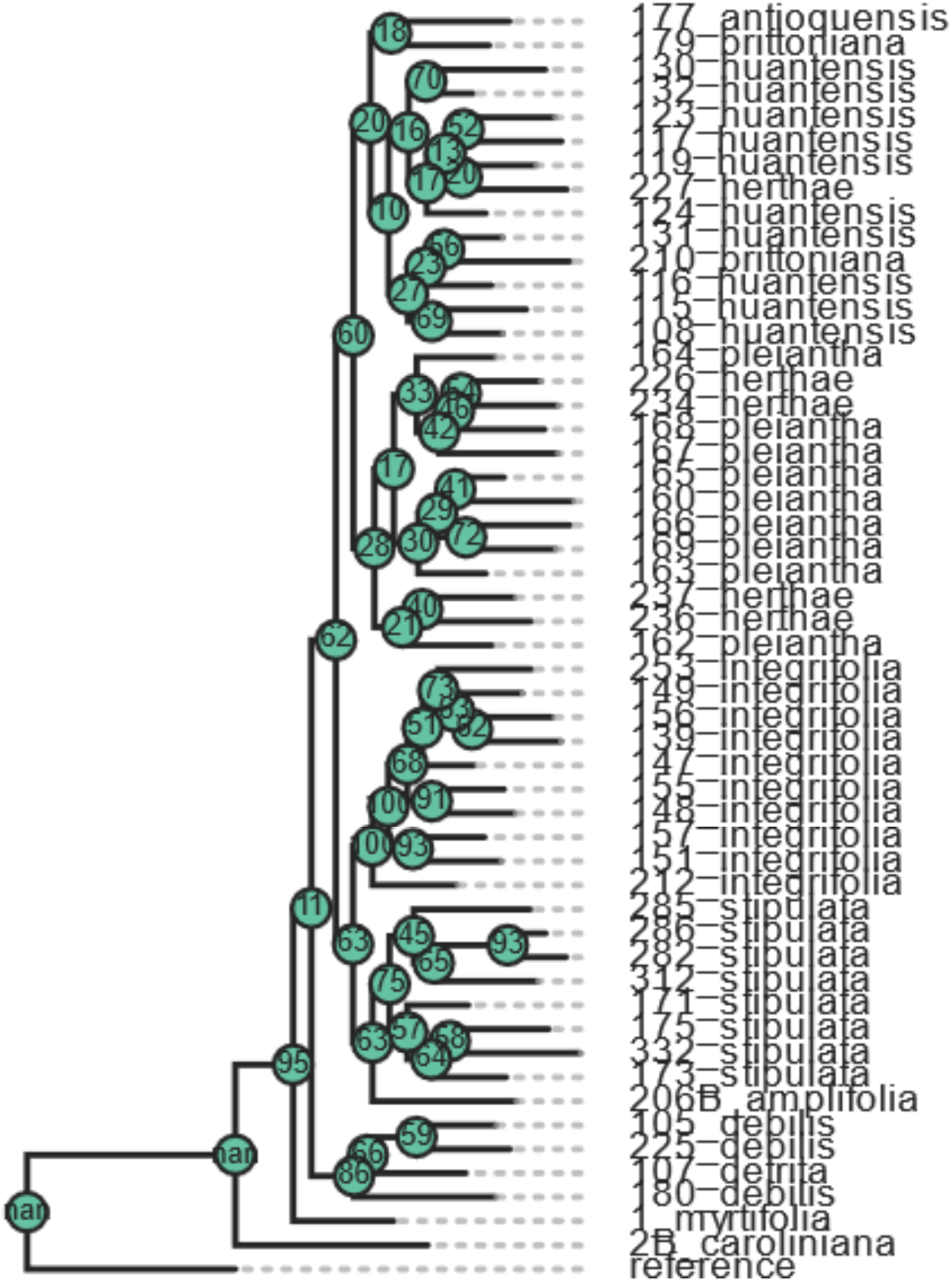
Maximum likelihood individual-level phylogeny reconstructed in RAxML. Individuals were selected to include those with at least 1000 loci per sample and up to 10 individuals per species. Sites were filtered to only include those with at least 25% coverage across samples. Tree was rooted to the reference (*P. persica*) and nodes show bootstrap support value.

**Figure S2.**
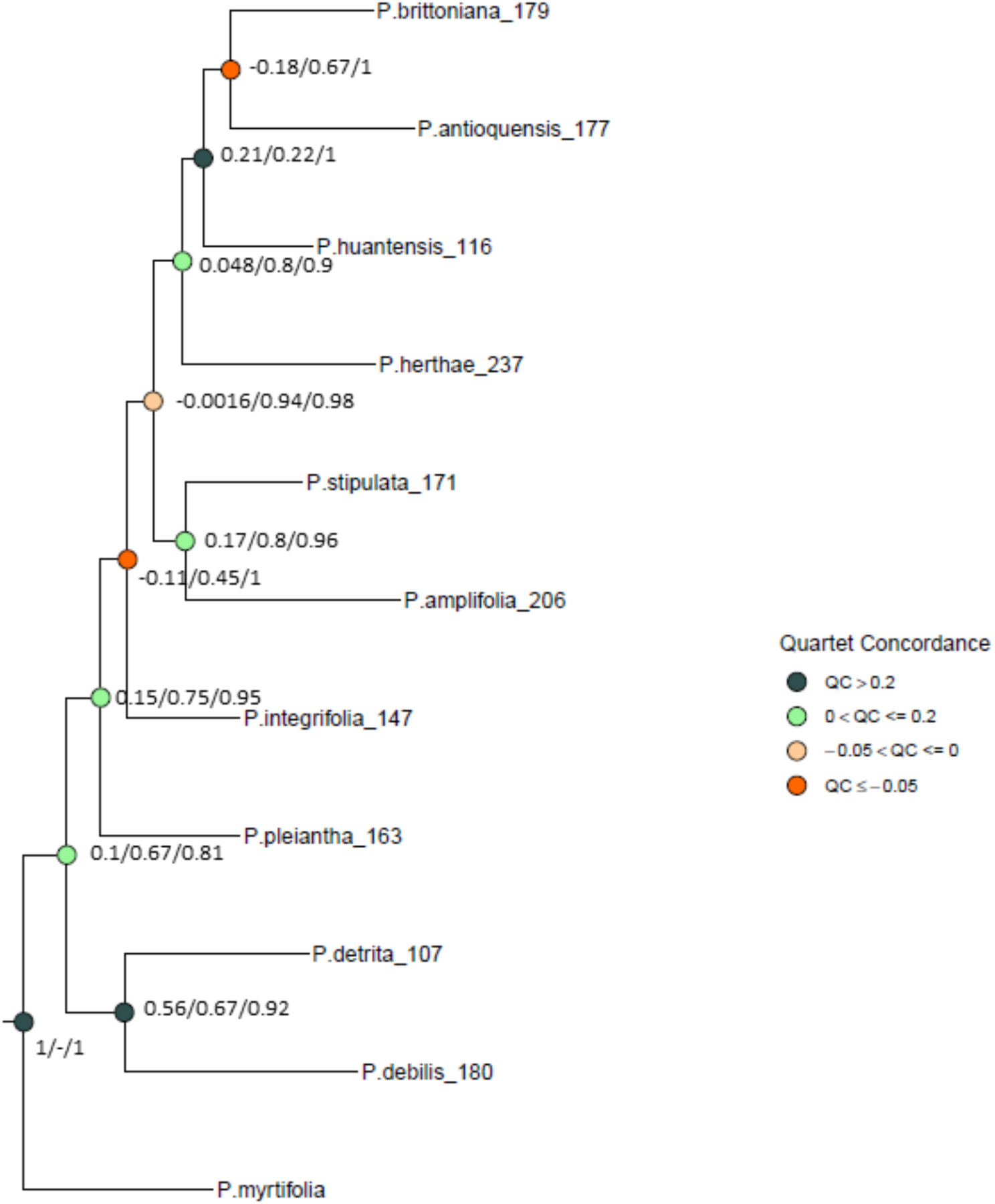
Quartet sampling results with quartet concordance (QC)/quartet differential (QD)/quartet informativeness (QI) indicated next to nodes. Nodes are colored according to QC scores, indicating how often the concordant quartet is inferred over both discordant quartets. Outgroup samples *P. persica* and *P. caroliniana* are removed for clarity. Figure created with https://github.com/ShuiyinLIU/QS_visualization.

**Figure S3.**
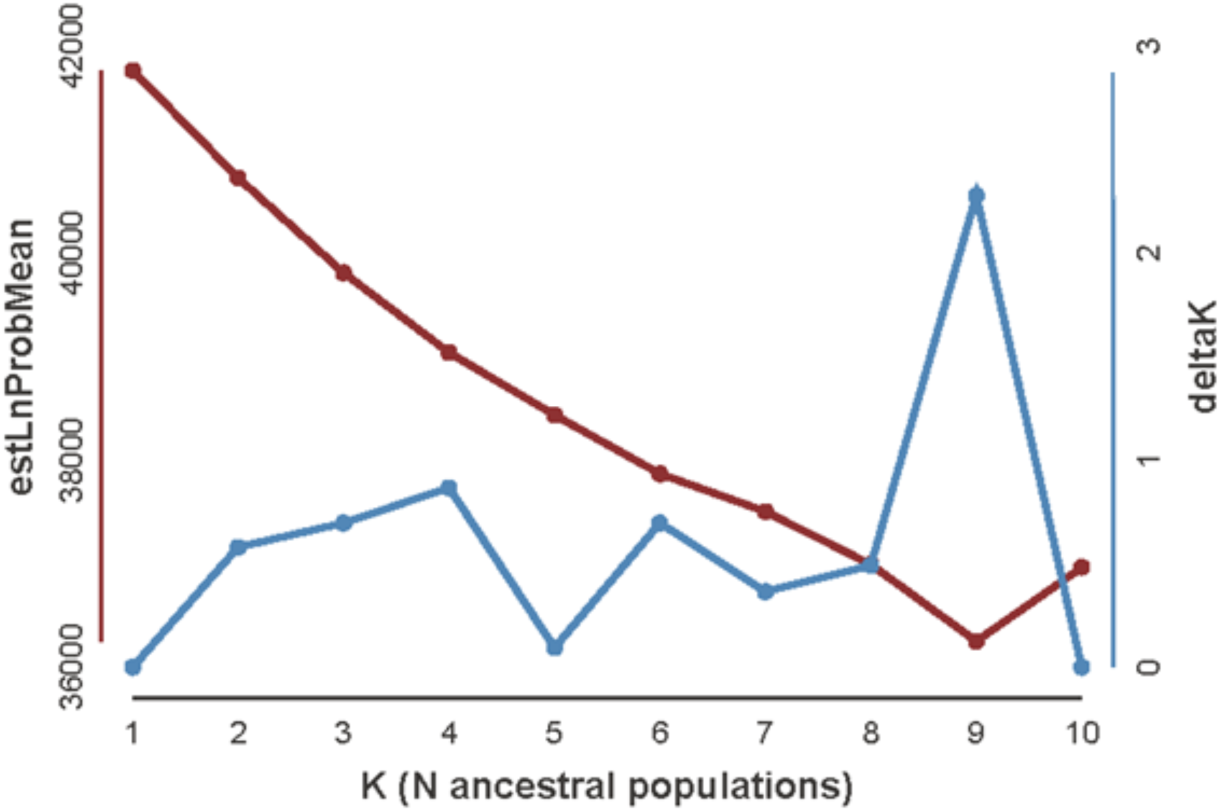
Plot of estimators used to select the optimal number of genetic clusters (K) in the STRUCTURE analysis. Estimated log mean probability assesses how well the model with a particular K explains the genetic data. As the log-likelihood values are negative, the best-fit K-value is the least negative. Delta K estimates the rate of change in the log probability of data between successive K values. The K value corresponding to the highest delta K and estimated log mean probability (K=9) was considered the best estimate of the number of subpopulations in this dataset.

